# Mode-based morphometry: A multiscale approach to mapping human neuroanatomy

**DOI:** 10.1101/2023.02.26.529328

**Authors:** Trang Cao, James C. Pang, Ashlea Segal, Yu-Chi Chen, Kevin M. Aquino, Michael Breakspear, Alex Fornito

## Abstract

Voxel-based morphometry (VBM) and surface-based morphometry (SBM) are two widely used neuroimaging techniques for investigating brain anatomy. These techniques rely on statistical inferences at individual points (voxels or vertices), clusters of points, or a priori regions-of-interest. They are powerful tools for describing brain anatomy, but offer little insights into the generative processes that shape a particular set of findings. Moreover, they are restricted to a single spatial resolution scale, precluding the opportunity to distinguish anatomical variations that are expressed across multiple scales. Drawing on concepts from classical physics, here we develop an approach, called mode-based morphometry (MBM), that can describe any empirical map of anatomical variations in terms of the fundamental, resonant modes––eigenmodes––of brain anatomy, each tied to a specific spatial scale. Hence, MBM naturally yields a multiscale characterization of the empirical map, affording new opportunities for investigating the spatial frequency content of neuroanatomical variability. Using simulated and empirical data, we show that the validity and reliability of MBM are either comparable or superior to classical vertex-based SBM for capturing differences in cortical thickness maps between two experimental groups. Our approach thus offers a robust, accurate, and informative method for characterizing empirical maps of neuroanatomical variability that can be directly linked to a generative physical process.

## 1. Introduction

Voxel-based morphometry (VBM) [1] and surface-based morphometry (SBM) [2] are the most commonly used techniques for studying neuroanatomical variations with magnetic resonance imaging (MRI). They are used to find associations between morphometric quantities (e.g., cortical thickness, surface area, grey matter volume) and diverse sociodemographic [3, 4, 5], behavioral [6, 7, 8], and clinical [9] variables. Throughout this article, we will focus principally on applications that examine neuroanatomical differences between two experimental groups (e.g., males and females, patients and controls), but our arguments and techniques easily generalize to cross-sectional analyses of inter-individual variability (e.g., brain-wide association analyses [10]).

The typical procedure in VBM- and SBM-based analyses of group differences involves fitting a statistical model at each individual voxel or vertex, respectively, and then applying some statistical threshold for inferring the spatial location(s) of significant mean differences [11, 12, 13]. In some cases, voxels/vertices are aggregated into anatomical regions-of-interest (ROIs) defined using an a priori atlas [14]. These approaches have been successful in describing where the differences are located in the brain, but offer no insights into the fundamental constraints that may have shaped those differences. Moreover, these approaches rest on the assumption that anatomical differences (1) are highly localized; (2) arise independently of each other; and (3) are confined to the spatial scale defined by the measurement resolution (i.e., local collections of voxels/vertices or atlas-based ROIs). These assumptions interact with the statistical thresholding procedure in a way that can obscure spatially extended patterns underlying the data (for an example, see Fig. S15 in ref. [15]). Moreover, the application of different statistical thresholds, derived using different methods, contributes to inconsistencies in findings reported in the literature [13, 16, 17, 18, 19]. Hence, a robust framework is needed that does not necessarily rely on a predefined threshold and can examine anatomical differences at multiple scales.

In diverse areas of physics and engineering, analysis at multiple scales can be obtained as the structure of a system can be comprehensively described by its structural eigenmodes, which are also referred to as modes, eigen-functions, eigenvectors, or harmonics [20, 21, 22]; we will use these terms interchangeably in the text. Eigenmodes correspond to the natural, resonant modes of the system and represent an orthogonal basis set that can describe any spatial pattern expressed by the system, much like the basis set of sines and cosines used to understand the frequency content of signals in Fourier analysis [20, 22]. Recent work has shown that eigenmodes derived either from a model of brain geometry, termed *geometric eigenmodes*, or from a graph-based model of the structural connectome based on diffusion MRI, termed *connectome eigenmodes*, can be used as a basis set for reconstructing diverse aspects of brain activity [23, 24, 25, 26, 27, 28, 15, 29, 30, 31, 32], for quantifying structure-function coupling in the brain [33, 34, 35], and for understanding atrophy patterns in neurodegeneration [36, 37, 38] and other conditions [39, 40]. In each of these cases, empirical spatial brain maps can be viewed as resulting from the preferential involvement, or excitation, of specific resonant modes of brain structure, thus offering insights into the generative physical mechanisms that shape the observed spatial pattern.

A further advantage of a mode-based approach is that, as in Fourier analyses, the modes are ordered by their topological (connectome eigenmodes) or spatial (geometric eigenmodes) frequency, offering a spectral decomposition of the data that provides insights into its multiscale organization. Indeed, recent work using geometric eigenmodes suggests that virtually all task-evoked activation maps obtained with functional MRI result from excitations of low-frequency modes spanning wavelengths > 60 mm [15]. Other work indicates that individual differences in brain anatomy are most salient at relatively coarse scales, spanning wavelengths ⩾37 mm [41]. Such findings challenge classical approaches to brain mapping that focus on identifying focal effects in isolated brain regions and underscore the benefits of a spatially-informed, multiscale approach.

Here, we leverage the advantages offered by a mode-based view of the brain to develop a new approach, called mode-based morphometry (MBM), for mapping anatomical differences between groups. This approach involves modeling such differences as a linear combination of eigenmodes and performing statistical inference on the modes, rather than on individual voxels or vertices. In this work, we use the geometric eigenmodes of the cortex, given their superior performance in explaining functional data [15], but note that our method can be adapted for use with other eigenmodes (e.g., connectome eigenmodes) as required. We focus on a surface-based analysis of cortical thickness (CT) differences, but other morphometric measures (e.g., surface area, volume, etc.) can easily be analyzed with the same method. We develop a framework for simulating CT maps with known ground truth and use them to compare the accuracy and reliability (under resampling) of MBM with classical vertex-based SBM. We then evaluate the consistency of the findings obtained by the two techniques as applied to empirical data through the analyses of (i) CT differences between sexes and (ii) multi-site measures of CT differences between healthy controls and patients with Alzheimer’s disease. In both simulated and empirical scenarios, our results indicate that MBM offers comparable, or superior, validity and consistency compared to SBM, while offering a direct insight into the generative mechanisms and multiscale characteristics of the data.

## 2. Methods

We begin by describing the details of the SBM and MBM approaches. We then describe our methods for evaluating the accuracy and consistency of the two approaches using simulated and empirical data.

### 2.1. Surface-based morphometry (SBM)

Traditional SBM compares neuroanatomical variations between two groups of interest using a general linear model (GLM) (Fig. 1a) given by

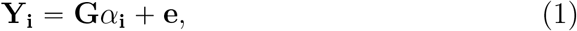

where **Y**_**i**_ is the data vector (*L* × 1) of *L* measurements at vertex *i* of the cortical surface, **G** is the design matrix (*L* × *P*) of effect variables, *α*_*i*_ is the parameter vector (*P* × 1), and **e** is an error vector (*L* × 1). Elements of *α*_*i*_ control the contribution of the corresponding effect columns of **G** to each vertex, **Y**_**i**_. Here, the GLM parameterizes the group differences as *t* statistics by defining a design matrix **G** and a contrast vector **c** as follows. Each element in a column of **G** has value 1 or 0 to indicate a measurement, i.e., a subject, belonging to a group or not. Each column of **G** controls the contribution of each group. The estimated values of *α*_*i*_ representing the mean of each group are multiplied by the contrast vector **c** defined by [1, −1] for calculating the *t*-statistic for each vertex (see [42, Appendix A] for more details).

**Figure 1.**
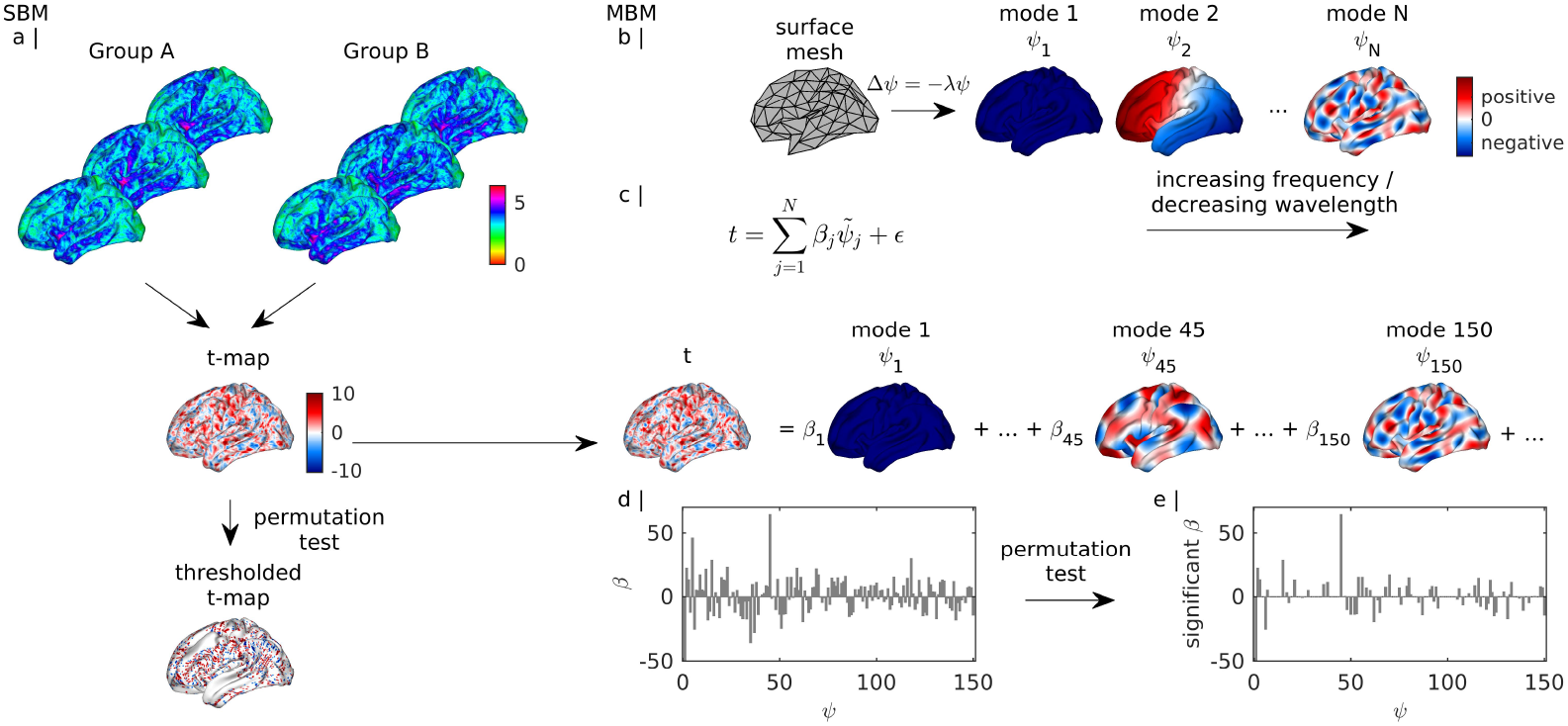
SBM and MBM analysis pipelines. (a) In SBM, a *t*-statistic is calculated independently at each vertex, quantifying point-wise group differences in CT. A thresholded *t*-map is derived by comparing the observed *t*-map and the distribution of null *t*-maps after permutation testing. (b) In MBM, eigenmodes are derived from a cortical surface mesh (solving Eq. (2)). The modes are ordered in increasing spatial frequency or decreasing spatial wavelength. Values in each mode are arbitrarily defined, with negative–zero–positive values in colored as blue–white–red. (c) An empirical *t*-map can be decomposed as a weighted sum of eigenmodes and errors using a GLM (Eq. 6), with weights given by *β*_*j*_. The set of *β*_*j*_ is called the *β* spectrum. (d) An example *β* spectrum with large *β*_45_ indicating a dominant contribution from mode 45. (e) An example of statistically significant *β*s derived by comparing the observed *β* spectrum and the distribution of null *β* spectra after permutation testing.

In this work, we performed non-parametric inference on the *t* statistics via permutation testing with 5000 iterations. At each iteration, we shuffled the group labels of all participants to create a *t*-map observed under random group assignment, resulting in an empirical null distribution at each vertex. We then used tail estimation on each null distribution [43] to calculate *p* values with arbitrarily high precision. In our analyses, we evaluated results with respect to (i) unthresholded differences; (ii) differences thresholded at *p* < 0.05 but uncorrected (*p*_unc_); and (iii) differences thresholded at *p* < 0.05 but corrected for false-discovery rate (FDR) (*p*_FDR_).

### 2.2. Mode-based morphometry (MBM)

MBM takes as input the unthresholded spatial map of *t* statistics quantifying group mean differences in CT and reconstructs that map using a basis set of structural eigenmodes of the brain. These eigenmodes can be defined using relevant anatomical properties of interest (e.g., geometry or connectome). The assumptions in MBM are that (1) the anatomical group differences reflect the superposition of different resonant modes of neuroanatomy that are preferentially expressed in one group over the other; and (2) these modes are orthogonal and intrinsic to the brain structure (see third paragraph of the Introduction for justification). Here, we focus on eigenmodes derived from the geometry of the cortex (geometric eigenmodes) because recent data [15] indicate that they offer a more accurate and parsimonious basis set for brain function and their derivation relies on a simpler processing pipeline. We provide further justification for this choice in the Discussion section.

The geometric eigenmodes are obtained by solving the eigendecomposition of the Laplace-Beltrami Operator (LBO), also known as the Helmholtz equation, on the cortical surface,

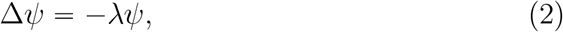

where Δ is the LBO and the solution *ψ* = {*ψ*_1_, *ψ*_2_, …} is the family of geometric eigenmodes with corresponding family of eigenvalues *λ* = {*λ*_1_, *λ*_2_, …}. The LBO captures the intrinsic geometry of the cortical manifold, which includes the curvature of the cortical surface in this case, and is generally defined as [44, 45, 15]

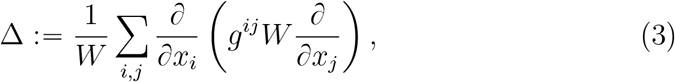

where *x*_*i*_, *x*_*j*_ are the local coordinates, 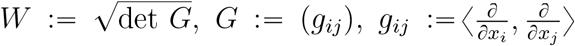 is an inner product, *g*^*ij*^ = (*g*_*ij*_)^−1^ is the inverse of *g*_*ij*_, and det denotes the determinant. *ψ* represents the set of spatially extended resonant modes, or eigenmodes, of the geometry. *λ* is related to the intrinsic spatial wavelengths of the modes, such that

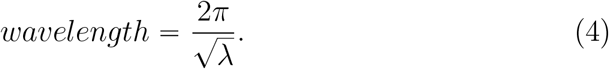

Eq. (4) is derived from the relationship?between eigenvalues *λ* and resonant or natural frequency *k* [45], with 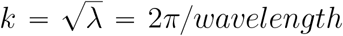. Furthermore, based on their wavelengths, the modes can be grouped into spatial scales by considering the special case of a sphere, which is known to be topologically similar to the cortical surface [46]. By solving Eq. (2) on the sphere, degenerate solutions exist, such that certain eigenmodes will have the same number of nodal lines and wavelengths, and they can be aggregated into an eigengroup *l*, similar to the angular momentum number in quantum physics. Each eigengroup comprises 2*l* + 1 eigenmodes, and its wavelength is [46, 41]

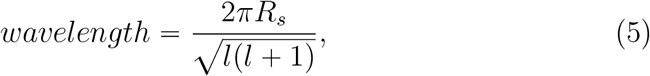

where *R*_*s*_ is the radius of the sphere (see Table S1 for an explicit list of wavelengths and eigenmode membership for each eigengroup on a sphere of *R*_*s*_ = 67 mm [41, 15]). Therefore, the eigenmodes of the cortical surface belonging to a given eigengroup have approximately the same spatial scale or wavelength given by Eq. (5).

The solutions of Eq. (2)—i.e., the geometric eigenmodes—correspond to the spatial part of the solutions of a wave equation describing the dynamics on a surface [47]. When applied to neuroanatomy, the eigenmodes represent an orthogonal basis set describing spatial patterns of the variations of cortical geometry at different spatial scales or wavelengths. The increasing order of the eigenvalues 0 ⩽ *λ*_1_ < *λ*_2_ < … corresponds to a decreasing order of the wavelengths. The first eigenvalue *λ*_1_ is approximately equal to zero with a wavelength that is very large compared to the size of the brain. Its corresponding eigenmode *ψ*_1_ is a constant function with no nodal lines and can be used to represent a mean across the brain.

We solved the eigenmodes of the cortical surface represented by a triangular mesh using the LaPy Python library [47, 48]. Here, we used a population-averaged template of the cortical midthickness surface in fsLR-32k CIFTI surface [49], comprising 32, 492 vertices in each hemisphere (Fig. 1b). We used a population-averaged surface to obtain a common set of eigenmodes for all participants, enabling easier comparisons. Note that individual-specific eigenmodes can also be derived from individual surfaces. However, small differences in the geometry of the cortical surface can alter the spatial patterns of the eigenmodes, which makes comparison between individual-specific modes challenging, particularly for short wavelengths or more fine-grained modes [41]. Moreover, prior work indicates that relying on a single, canonical basis set of modes derived from a group template leads to a negligible difference in reconstructions of empirical data when compared to using individual-specific modes [15], suggesting that the group-derived modes provide a reasonable approximation for present purposes. Here, we performed analyses on the left hemisphere and thus used the modes derived from the left hemisphere.

We then used the eigenmodes to decompose the *t*-map of each hemisphere via a GLM given by (Fig. 1c)

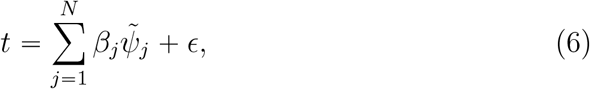

where *t* = [*t*_1_, *t*_2_, …, *t*_*i*_, …] is the *t*-map, *i* represents a vertex, 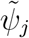 is the normalized *j*th eigenmode, *β*_*j*_ is the coefficient quantifying the contribution of 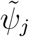 to the *t*-map, *N* is the number of modes used, and *ϵ* is a constant vector (error vector). Note that *β*_*j*_ is distinct from the eigenvalue *λ*_*j*_, with the former corresponding to the contribution of eigenmode *j* to the *t*-map and the latter corresponding to the wavelength of eigenmode *j*. In this work, we considered *N* = 150 modes spanning wavelengths ⩾34 mm, given recent work showing that the most salient aspects of brain function reside at spatial wavelengths longer than 34 mm [15, 41]. However, our results do not change with other choices of *N* (see Fig. S1). We normalized the eigenfunction *ψ*_*j*_ by dividing it with its Euclidean norm to ensure orthonormality of the basis set.

We call the set of *β*_*j*_ coefficients the *β* spectrum, which encodes the contribution of each eigenmode at a specific spatial scale, i.e., each frequency component, to the neuroanatomical property being investigated. In the present application, the investigated property is a *t*-statistic map quantifying group differences in CT. Thus, for example, if *β*_45_ is large, the spatial pattern defined by mode 45 dominates group differences in CT (Figs. 1c and d).

We performed statistical inference on the *β* estimates using permutation testing. To this end, we reconstructed null *t*-maps obtained after shuffling the group labels, as described in Section 2.1 for the SBM analysis, yielding an empirical distribution of null *β* spectra. We then used tail approximation to estimate *p*-values for each observed *β* value in the spectrum [43]. This procedure thus allows us to determine which specific modes make a significant contribution to the observed CT differences between groups. In this way, MBM performs inference at the level of the modes of brain structure rather than individual voxels. The analysis thus determines the specific modes that make a statistically significant contribution to the empirical spatial map. As in the SBM analysis, we considered unthresholded spectra and those thresholded at *p*_unc_ < 0.05 and *p*_FDR_ < 0.05 (Fig. 1e).

### 2.3. Validations using simulated data

One challenge in evaluating the validity of any new brain mapping technique is the lack of a ground truth for most applications. Here, we developed a framework for simulating group differences in CT with a known ground truth to compare the performance of SBM and MBM.

#### 2.3.1. Simulation Framework

We simulated two experimental groups based on empirical CT maps using the simple model defined in Eq. (7). We simulated the CT map, **y**, of a subject in each group by combining three elements: (1) a group-specific common phenotype map (**M**_*C*_), which represents a structured CT pattern, i.e., a ground truth, that is common to all members of a group, thus justifying their assignment to the same group; (2) an individual-specific structured noise map (**M**_*S*_), which represents structured CT variations that are specific to each individual; and (3) Gaussian noise (**M**_*G*_), which represents measurement error. **M**_*S*_ reflects between-subject variance, which can be considered structured noise when compared with the group means. It is structured in the sense that it is autcorrelated. For simplicity, we assumed that the combination of these three elements was linear, such that the model for **y** was

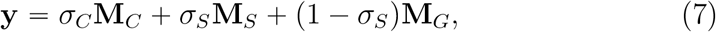

where *σ*_*C*_ (0 ⩽ *σ*_*C*_ ⩽ 1) is a free parameter that controls the contribution of the group-specific CT phenotype, **M**_*C*_, and *σ*_*S*_ (0 ⩽ *σ*_*S*_ ⩽ 1) is a free parameter that controls the relative contributions of structured noise, **M**_*S*_, and Gaussian noise, **M**_*G*_ (Fig. 2a). When *σ*_*C*_ = 0, the generated maps only contain noise, and when *σ*_*C*_ = 1, the generated maps have equal weightings between the group-specific map, representing the ground-truth phenotype, and noise, including structured and Gaussian noise. When *σ*_*S*_ = 0, the generated maps are only affected by Gaussian noise, and when *σ*_*S*_ = 1, the generated maps are only affected by individual-specific structured noise. Note that the range of *σ*_*C*_ and *σ*_*S*_ were chosen for simplicity to capture the relation between **M**_*C*_, **M**_*S*_, and **M**_*G*_.

**Figure 2.**
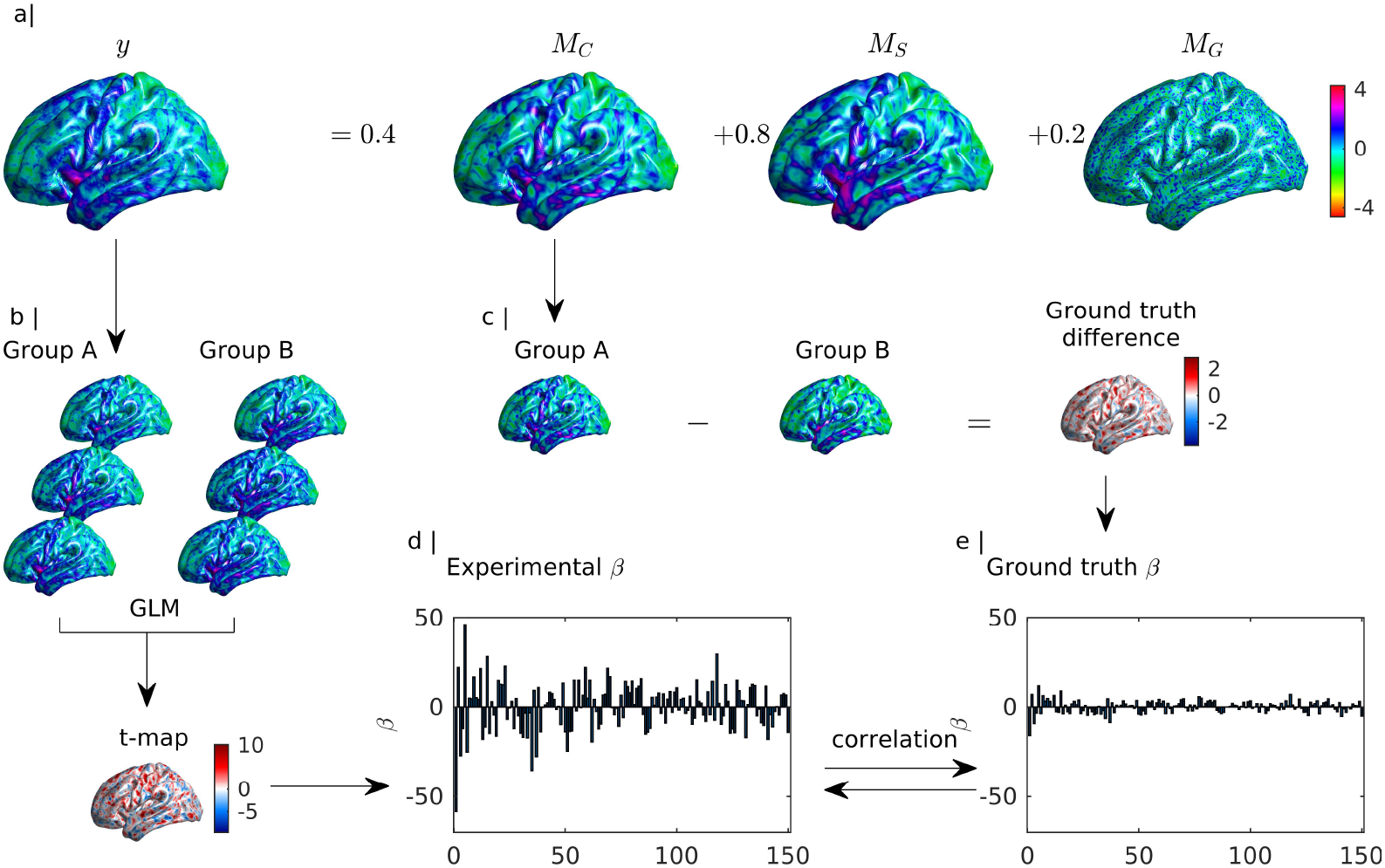
Framework for ground-truth simulations. (a) We generate a CT map using the model in Eq. (7). Here, we show an example with *σ*_*C*_ = 0.4 and *σ*_*S*_ = 0.8. (b) Using the simulated CT maps for groups *A* and *B*, we estimate a *t*-map and its corresponding *β* spectrum (d). (c) The ground-truth (GT) difference map is given by the subtraction of 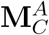 and 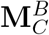, from which the ground-truth *β* spectrum (e) is obtained.

To specify two groups *A* and *B*, we include the superscripts *A* and *B* to **M**_*S*_ and **M**_*C*_; for instance 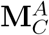 denotes the common group CT phenotype map of a subject in group *A*. Under this model, classical inference on group differences is the same as asking whether there is a difference in 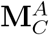 and 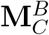 that can be reliably detected over and above the variability associated with **M**_*S*_ and **M**_*G*_.

We used 25 subjects for each group, unless otherwise stated, to mimic sample sizes historically used in the literature, although our general conclusions are not affected by this choice (see Fig. S2). To ensure that the generated maps have the same spatial structure as empirical data, we sampled maps **M**_*C*_ and **M**_*S*_ without replacement from 339 CT maps in the Human Connectome Project (HCP) data described in Sec. 2.4.1 [50]. Note that **M**_*C*_ corresponds to the same spatial map for all participants within a group, whereas a different **M**_*S*_ was selected for each individual within a group. We embedded actual empirical data within the simulations in this way to preserve the real structure and properties of CT maps as much as possible.

For each **y** map, we subtracted the minimum negative value across all **y** (both groups) to remove negative values caused by the addition of Gaussian noise to the CT maps. Given Eq. (1), this offset did not affect the *t*-map comparing group mean CT differences, as the same constant value was subtracted to the two groups.

For each pair of parameters *σ*_*C*_ and *σ*_*S*_, we ran 100 experiments with fixed ground-truths 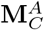 and 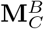. We chose 100 experiments to balance the computational cost and the reliability of the resulting performance metrics. Figure S3 confirms that our findings are not sensitive to the selection of different ground truths. From the generated CT maps of the two groups, the *t*-map and *β* spectrum of each experiment were computed as described in Sections 2.1 and 2.2, respectively (Figs. 2b and d).

The parameters *σ*_*C*_ and *σ*_*S*_ allow us to evaluate the sensitivity of both SBM and MBM to variations in the level of real phenotype and structured and Gaussian noise. To determine the most plausible parameter regime that yields the most realistic estimates of empirical data, we compared the spatial variograms of the empirical and simulated CT maps under different parameter combinations (Fig. 3). A variogram provides a measure of spatial dependence, or spatial autocorrelation, of a random field, defined as the variance of the difference between field values at two locations across realizations of the field. We used the Python package BrainSMASH [51] to calculate the variograms. For the empirical data, we calculated the mean and variance of the variograms of the 339 CT maps from the HCP data (see Section 2.4.1). For each pair of parameters *σ*_*C*_ and *σ*_*S*_ in the simulations, we generated 500 CT maps and calculated the mean and variance of the corresponding variograms. The slope of the variogram encodes the spatial structure of the data [51]. For example, a positive slope means that there is high autocorrelation in the data; i.e., variations at small distances are smaller than variations at large distances. A flat variogram means that variations are independent of distances. Similar slopes refer to the similarity of the spatial autocorrelations. Thus, to compare the slope of the mean of variograms of empirical maps and that of simulated maps, we removed the offset between their minimum values, so that the two variograms start at the same point (Fig. 3a). In particular, the offset was calculated from the mean variogram that has a higher minimum value. Note that removing the offset does not affect the structure of the variograms. We then calculated the norm distance between the emirical and simulated mean variograms after subtracting the minimum offset.

**Figure 3.**
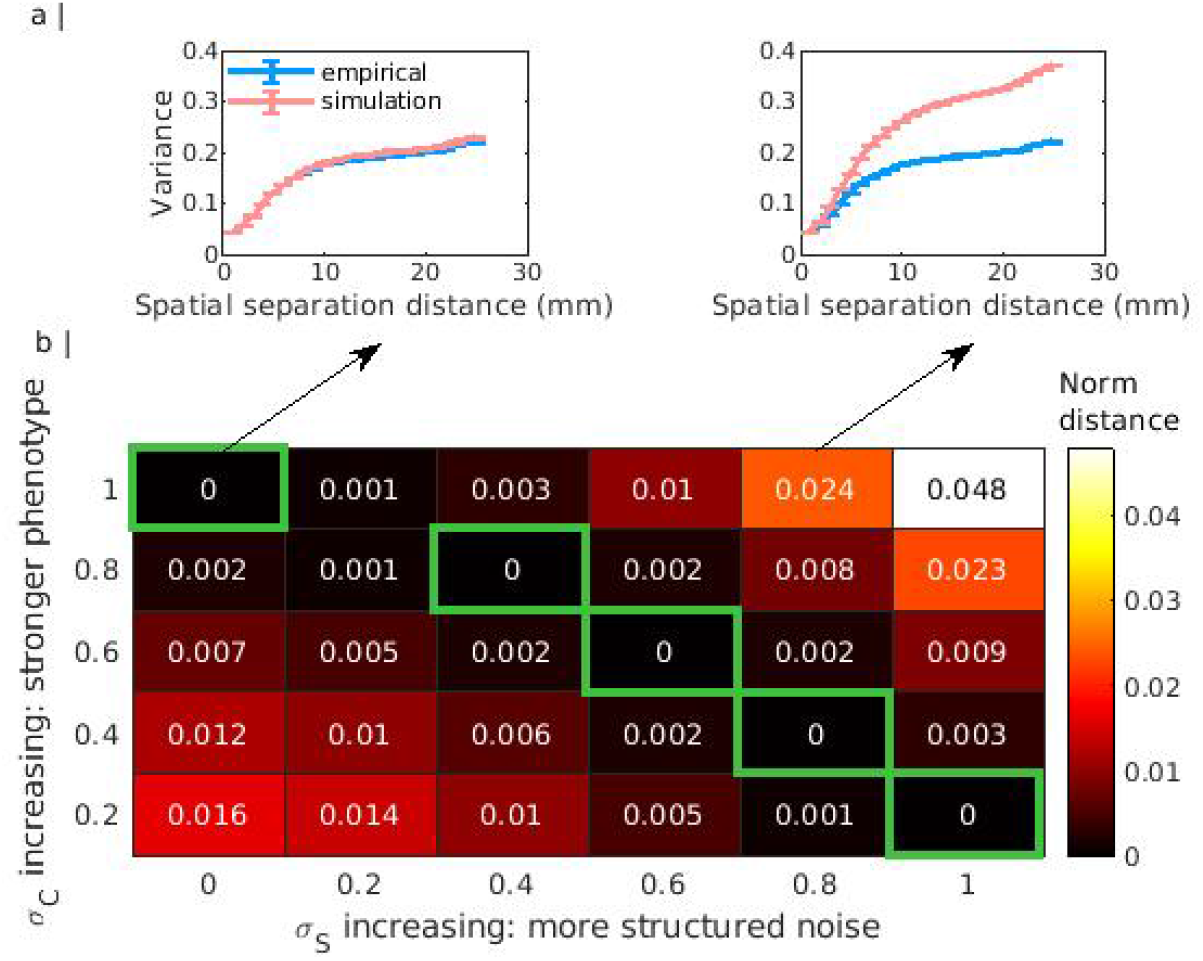
Comparing spatial variograms of empirical and simulated CT maps for different combinations of *σ*_*C*_ and *σ*_*S*_. (a) Two examples of mean variograms (after subtracting the minimum offset) of empirical and simulated maps for parameter pairs (*σ*_*C*_ = 1, *σ*_*S*_ = 0) and (*σ*_*C*_ = 1, *σ*_*S*_ = 0.8). The error bars (vertical bars) show the variance of the variograms. (b) Norm distances between the empirical and simulated mean variograms (after subtracting the minimum offset) for combinations of *σ*_*C*_ and *σ*_*S*_. The green boxes highlight the realistic regimes where the generated maps have a similar spatial structure as the empirical data (norm distance ≈ 0).

Figure 3b shows the norm distances for combinations of *σ*_*C*_ and *σ*_*S*_. Figure 3a shows two examples of the empirical and simualted mean variograms (after subtracting the minimum offset) at parameter pairs (*σ*_*C*_ = 1, *σ*_*S*_ = 0) and (*σ*_*C*_ = 1, *σ*_*S*_ = 0.8) (see Fig. S4 for the mean variograms of all *σ*_*C*_ and *σ*_*S*_ in the norm distance table). The green boxes in Fig. 3b highlight the realistic parameter regimes where the generated maps have a similar spatial structure to the empirical data; i.e., the norm distances are close to zero. We will highlight these realistic regimes in the Results section and examine the effects of parameter choices in these realistic regimes.

#### 2.3.2. Performance evaluation

We compared the performance of SBM and MBM in terms of accuracy and consistency. We defined accuracy as the ability of each method to accurately capture the ground-truth group difference in the simulations. These ground-truth difference maps were obtained by subtracting the 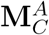 and 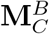 maps (Fig. 2c). The accuracy of SBM was quantified as the product-moment correlation between this ground truth and the *t*-map obtained through classical analysis (see Section 2.1). Similarly, the accuracy of MBM was quantified as the product-moment correlation between the *β* spectrum (see Section 2.2) of the ground-truth difference map (Fig. 2e) and the *β* spectrum obtained for each experiment (Fig. 2d).

We defined consistency as the ability of each method to obtain consistent findings in the face of sampling variability across repeated experiments. Therefore, we compared the distributions of pairwise correlations between *t*-maps of 100 experiments for SBM and pairwise correlations between *β* spectra of 100 experiments for MBM. When considering thresholded results, we used binary correlations [52, 53]. We present unthresholded and uncorrected thresholded results at *p*_unc_ ⩽ 0.05 in the main text, and FDR-corrected results in Supplementary Fig. S5. The general conclusions are consistent for both uncorrected and corrected results. We also evaluated the performance of SBM and MBM on CT maps spatially smoothed with surface-based kernel sizes of 10 mm, 20 mm, or 30 mm full-width at half-maximum (FWHM) to match with the common smoothing practice of SBM. The Connectome Workbench software [54] was used to smooth the CT maps.

### 2.4. Validations using empirical data

The simulated data offer an opportunity to compare the accuracy of the proposed MBM method vs SBM with respect to a known ground truth. Ideally, we could also perform a similar analysis on empirical data; however, ground truth is difficult to ascertain for such applications. Hence, we can only assess the consistency of MBM and SBM in analyzing empirical data, provided that we have multiple runs of comparable experiments. Here, we evaluated consistency with respect to two different group comparisons and empirical datasets, as described below.

#### 2.4.1. Sex differences

The first empirical validation focused on sex differences in CT maps from HCP [50]. In particular, we analyzed FreeSurfer-derived CT maps [55] from 339 unrelated healthy young adults (ages 22 to 35; 182 females; no siblings). This subsample corresponds to unrelated subjects from the HCP S900 release. The CT maps were spatially normalized to the fsLR-32k CIFTI surface using FreeSurfer [56, 57] and Connectome Workbench [54].

For each experiment, we subsampled 25 subjects from each group of females and males. We then computed the *t*-map and *β* spectrum for each experiment. We repeated this process 100 times, representing 100 experiments. We then compared the distributions of the correlations between *t*-maps and *β* spectra obtained for each pair of experiments, as in the consistency analysis using simulated data.

#### 2.4.2. Patient-control differences

The second empirical validation focused on patient vs control differences in CT maps from the Open Access Series of Imaging Studies (OASIS-3) [58]. We analyzed FreeSurfer-derived CT maps from 693 healthy individuals and individuals with Alzheimer’s disease (ages 42 to 95; 407 females). For subjects with multiple sessions, we only analyzed data from their last scan. The OASIS-3 data were acquired from one imaging center with three different 3T scanners and one 1.5T scanner. Hence, we treated data from each 3T scanner as a separate site and analyzed each site independently (we do not use data from the 1.5T scanner). The numbers of cognitively normal healthy controls (HC) and individuals with Alzheimer’s disease (AD) in each site are: site 1: 164 HC, 27 AD; site 2: 218 HC, 24 AD; and site 3: 142 HC, 118 AD. We excluded subjects with known history of active medicine-induced cognitive dysfunction, neurological diseases, seizure disorder, hydrocephalus, head injury, alcoholism, or Parkinson’s disease to minimize confounds.

Each of the three sites thus represents an independent experimental replication. We therefore evaluated the consistency of the findings obtained at each site using correlations between *t*-maps and *β* spectra between each pair of sites, as per the other analyses.

### 2.5. Analyzing the frequency content of group different maps

We used MBM to examine the frequency content of group differences in CT, e.g., the dominant scale of the group differences, in two ways. First, we assigned eigenmodes to distinct eigengroups, as outlined in Section 2.2, and quantified the fraction of significant modes in each group. Second, we evaluated the accuracy of mode-based reconstructions of the empirical CT difference map following incremental, sequential removal of modes according to spatial wavelength. Specifically, we removed modes starting from long-wavelength to short-wavelength modes (from the first mode to mode *i*) or starting from short-wavelength to long-wavelength modes (from the last mode to mode *i*), and evaluated the correlation between the empirical CT difference map and the map reconstructed using the remaining set of modes obtained after each removal.

## 3. Results

### 3.1. Simulated data

Figure 4a shows the mean correlations between *t*-maps and the ground-truth CT difference map obtained with SBM, hereafter referred to as SBM correlations. As expected, the SBM correlation increases as *σ*_*C*_ increases, which reflects a stronger contribution from the ground-truth CT pattern to the simulated CT map. Figure 4b shows a similar behavior for the mean correlations between the *β* spectra of *t*-maps and the ground-truth difference map obtained with MBM, hereafter referred to as MBM correlations. Critically, MBM has better performance for nearly all parameter combinations, including in the realistic parameter regimes denoted by the green boxes. Differences between SBM and MBM were particularly salient in regimes characterized by a weak phenotype contribution and high levels of Gaussian noise (i.e., low *σ*_*C*_ and low *σ*_*S*_). This effect likely reflects the low-pass spatial filtering effect of the MBM reconstruction, which renders it more robust to high-frequency noise than SBM. Thus, across a wide range of signal-to-noise ratios, MBM more accurately captures the ground-truth difference than SBM.

**Figure 4.**
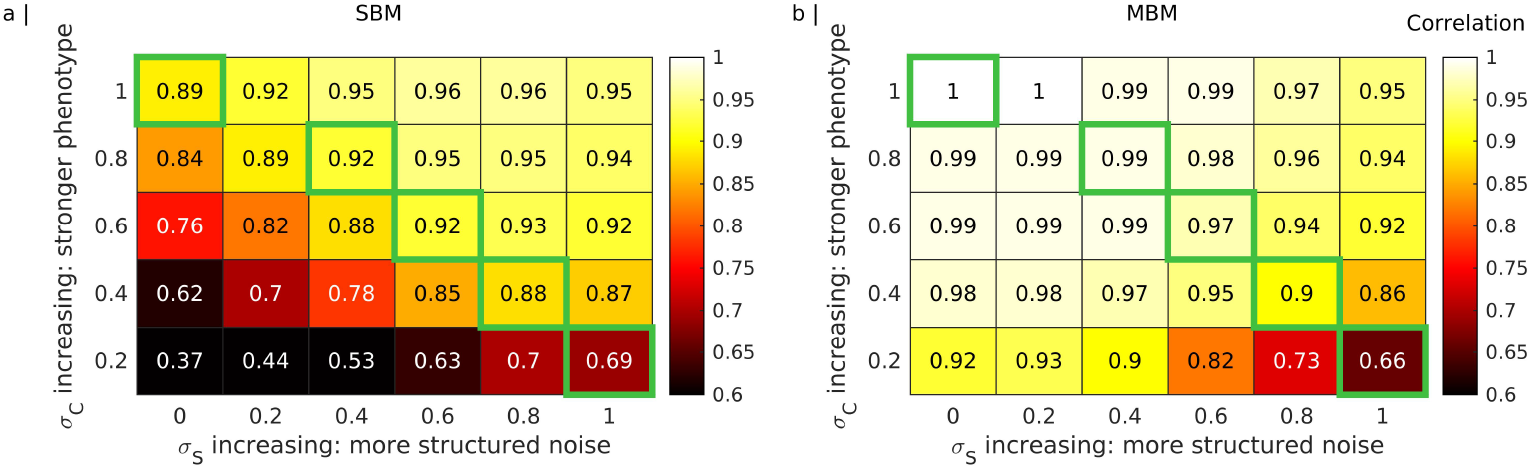
Accuracy of SBM and MBM with respect to ground-truth simulations for different combinations of *σ*_*C*_ and *σ*_*S*_. (a) Mean correlation between the *t*-map of an experiment and the ground-truth difference map. (b) Mean correlation between the *β* spectra of the *t*-map of an experiment and the ground-truth difference map. The green boxes highlight the realistic parameter regimes where the generated maps have a similar spatial structure as the empirical data, as shown in Fig. 3.

Figure 5 shows the results of our consistency analysis quantified in terms of the distributions of the SBM and MBM correlations, for both unthresholded and thresholded results, between each pair of 100 different experiments relying on the same ground truth. When *σ*_*C*_ increases, the distributions shift to the right and become narrower. Thus, when the contribution of the ground-truth difference is more strongly expressed in individual CT maps, both SBM and MBM become more consistent. Across most parameter combinations, including those in the realistic regimes, the distribution of MBM correlations shifts to the right of the SBM correlations for both unthresholded and thresholded results, indicating that MBM generally yields more consistent findings. The exception to this rule is in cases of a weak phenotype contribution and high levels of structured noise (i.e., low *σ*_*C*_ and high *σ*_*S*_), where the peaks of the SBM and MBM distributions converge but the latter show a wider variation. This result arises because MBM is particularly sensitive to structured spatial patterns extending over long wavelengths. MBM will thus have difficulty in reliably detecting a ground-truth difference in the presence of a high degree of structured noise such as, for instance, when individual variability swamps consistent patterns observed across different individuals belonging to the same group.

**Figure 5.**
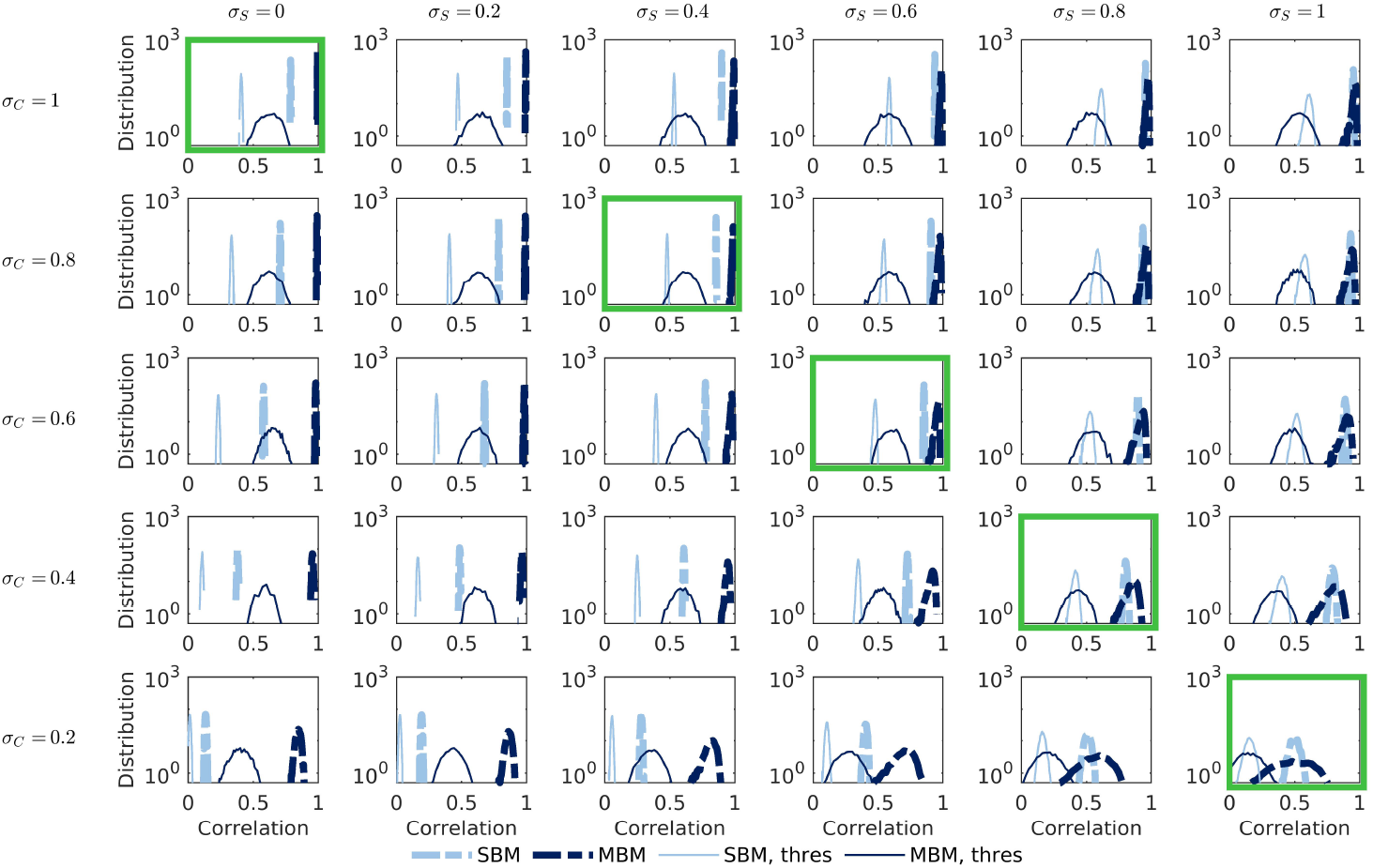
Distributions (in log scale) of correlations between pairs of 100 experiments for different combinations of *σ*_*C*_ and *σ*_*S*_. The panels show correlations between pairs of experimental *t*-maps (SBM), correlations between pairs of experimental *β* spectra (MBM), binary correlations between thresholded *t*-maps (SBM, thres), and binary correlations between statistically significant *β* spectra (MBM, thres). The green boxes highlight the realistic parameter regimes where the generated maps have a similar spatial structure as the empirical data, as shown in Fig. 3.

To further investigate this spatial filtering effect of MBM, we repeated the analyses after smoothing the CT data using surface-based smoothing kernels with FWHM of 10, 20, and 30 mm. Figure 6 shows the distributions of SBM and MBM correlations obtained for 5 pairs of *σ*_*C*_ and *σ*_*S*_ in realistic regimes, i.e., (*σ*_*C*_, *σ*_*S*_) = {(1, 0), (0.8, 0.4), (0.6), (0.6, 0.4), (0.8), (0.2, 1)}. When the smoothing kernel size increases, the SBM correlation distributions shift to the right and become broader, suggesting that smoothing improves SBM consistency on average, but large deviations from this average are also more common. On the other hand, the MBM correlation distributions are relatively unaffected by spatial smoothing. In general, smoothing only improves the consistency of SBM to a level commensurate with MBM applied to un-smoothed data. This result implies that MBM offers similar advantages to smoothing without having to choose an arbitrary kernel size and shape.

**Figure 6.**
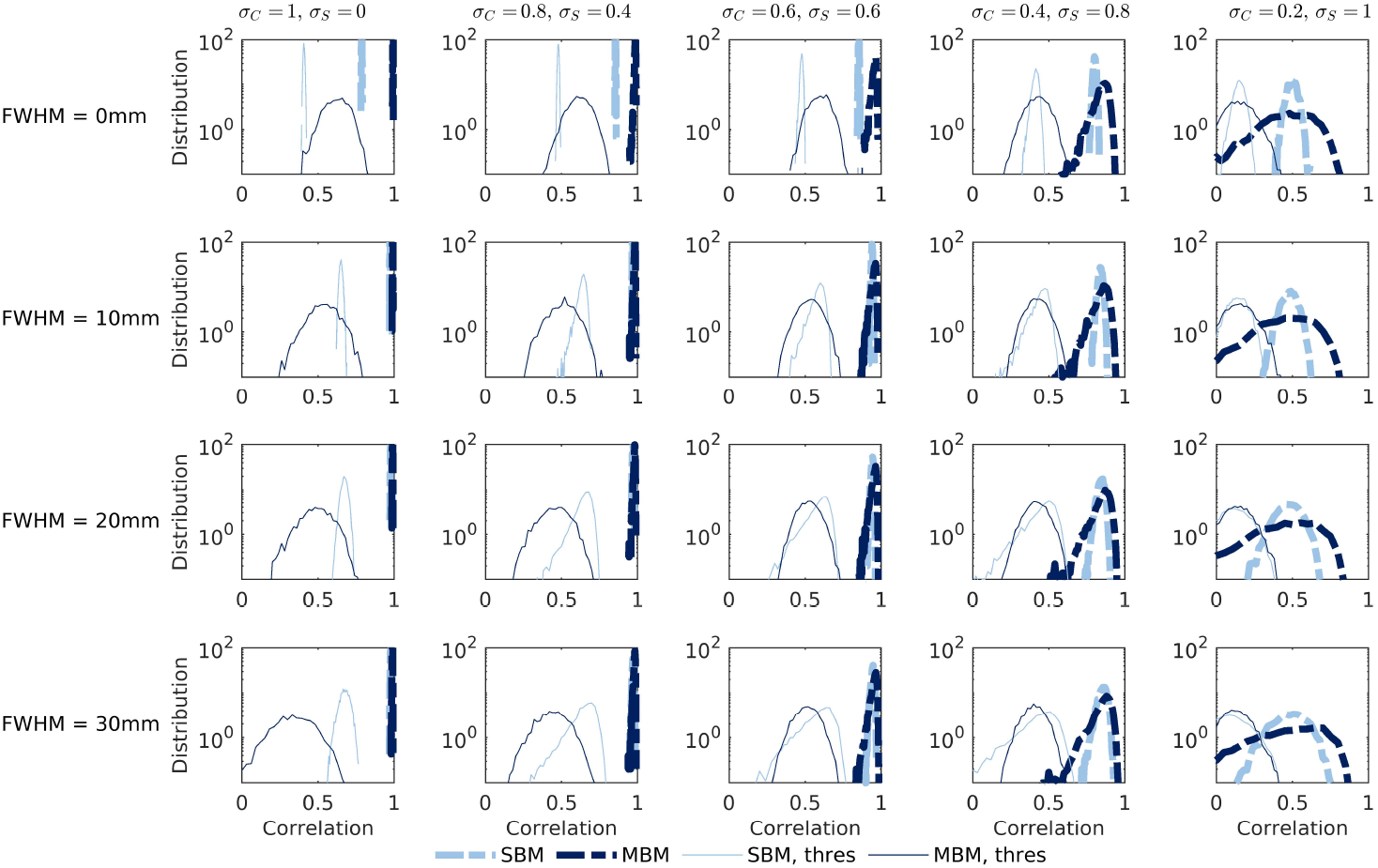
Distribution (in log scale) of pairwise correlations between experiments in the realistic parameter regimes and for different smoothing kernels. The panels show correlations for experimental *t*-maps (SBM), correlations for experimental *β* spectra (MBM), binary correlations for thresholded *t*-maps (SBM, thres), and binary correlations for statistically significant *β* spectra (MBM, thres).

### 3.2. Empirical results

Having demonstrated that MBM yields more accurate and consistent results than SBM in simulated data, we now evaluate the performance of the two methods in actual empirical data. To reiterate, since there is no clear ground truth in such cases, we primarily focus on evaluating the consistency, under sampling variability, of results obtained with both methods.

#### 3.2.1. Sex differences

Figure 7 shows the SBM and MBM analyses of CT differences between sexes. Figures 7a and b show the *t*-map and thresholded *t*-map in a typical SBM. Figure 7c shows the *β* spectrum obtained from MBM. The *β* spectrum describes the influence of multiscale patterns, i.e., modes, on the CT differences. As shown in Fig. 7c, CT differences between sexes are influenced more by long-wavelength, i.e., low-frequency, modes than short-wavelength, i.e., high-frequency, modes. For example, *β*s of modes 1 to 50 have higher absolute amplitudes compared to those of modes 100 to 150. The significant modes with their *β*s (shown in green in Fig. 7c) are combined to show the significant pattern of the CT difference between sexes in Fig. 7d. The six most influential modes, i.e., significant modes with largest absolute *β* values, are shown in Fig. 7e. Figure 7f shows the smoothed *t*-map at FWHM=30 mm, which is different from Fig. 7d, demonstrating that the complex pattern of CT difference extracted by MBM cannot be trivially obtained by simply smoothing the *t*-map.

**Figure 7.**
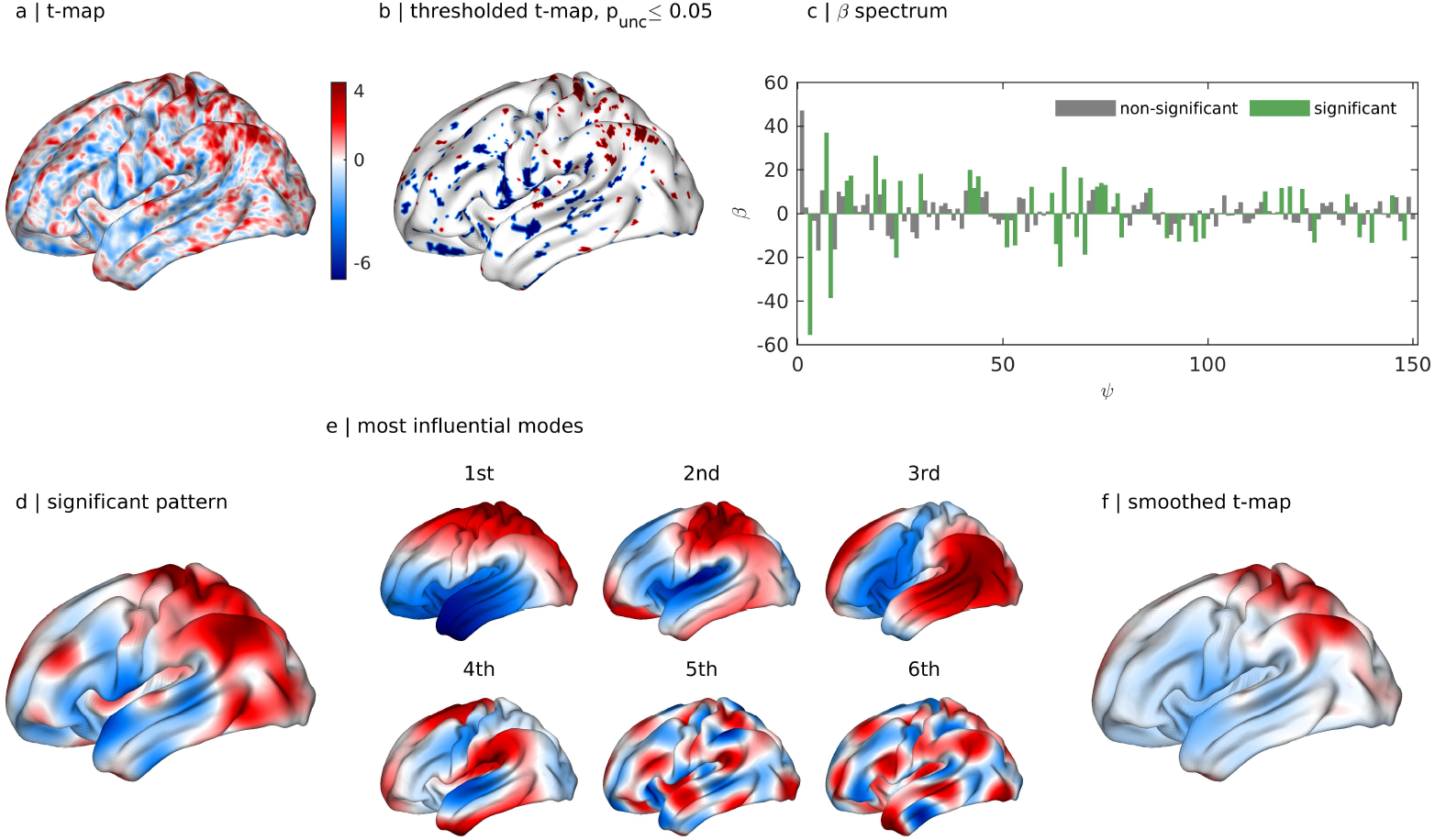
SBM and MBM analyses of CT differences between sexes. (a) Unthresholded *t*-map. (b) Thresholded *t*-map at *p*_unc_ ⩽ 0.05. Red and blue denote significantly thicker CT in females and males, respectively. (c) *β* spectrum of the unthresholded *t*-map. The *β*’s of the significant modes, obtained via permutation testing at *p*_unc_ ⩽ 0.05, are colored green. (d) Significant pattern obtained by combining the significant modes weighted by their *β*’s. (e) Six most influential modes, i.e., significant modes with largest absolute *β* values. The signs of modes with negative *β*’s were flipped to better visualize the similarity between the modes and the significant patterns. The number denotes the order of influence, not the mode index. (f) Smoothed *t*-map at FWHM = 30 mm.

Figure 8a shows that MBM yields more consistent results than SBM when considering sex differences in the HCP data under resampling. Figures 8b-d indicate that applying progressively larger smoothing kernels leads to a convergence in the consistency of both SBM and MBM, but SBM never displays clearly superior performance. These results align with our simulation findings. In fact, the wide distribution tails observed with MBM, particularly at higher smoothing kernels, suggest that the empirical sex difference maps studied here are characterized by a relatively weak true phenotype contribution (i.e., a reliable, characteristic sex difference) and a strong contribution of structured noise; in other words, the data appear to correspond to a low *σ*_*C*_, high *σ*_*S*_ regime.

**Figure 8.**
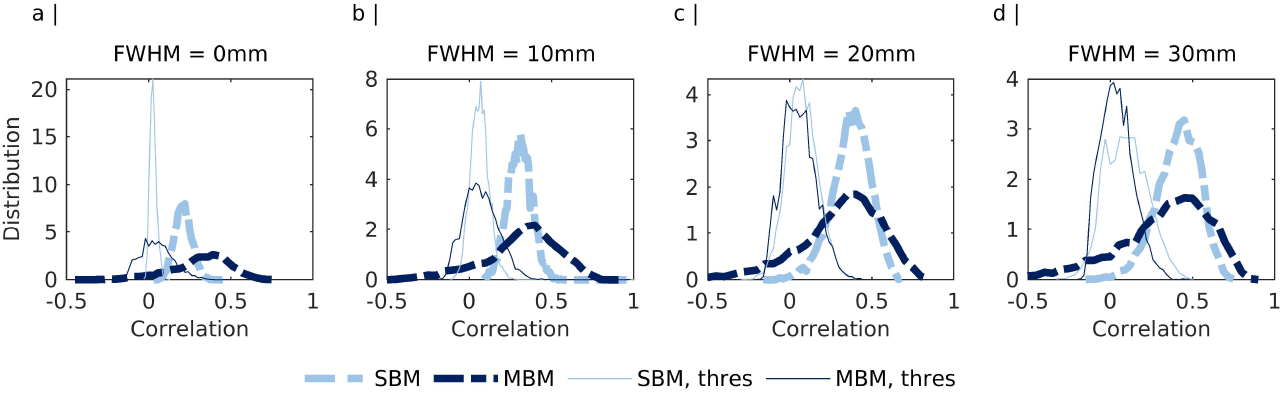
Distribution of pairwise correlations between experiments for different smoothing kernels in the empirical analysis of sex differences under resampling. The panels show correlations for experimental *t*-maps (SBM), correlations for experimental *β* spectra (MBM), binary correlations for thresholded *t*-maps (SBM, thres), and binary correlations for statistically significant *β* spectra (MBM, thres).

#### 3.2.2. Alzheimer’s disease

Figures 9a and b show the *t*-maps and thresholded *t*-maps of CT differences between patients with Alzheimer’s disease and healthy controls across three sites. Figures 9c, d, and e show the results obtained from MBM, including the *β* spectra, reconstructions using the significant patterns of the CT difference, and the six most influential modes. The reconstructions using significant modes show a consistent pattern of differences across the sites that is not directly apparent when inspecting the thresholded *t*-maps obtained with SBM. Accordingly, the six most influential modes show consistency across the datasets and all sites include the first mode, implying consistent global differences in CT between cases and controls.

**Figure 9.**
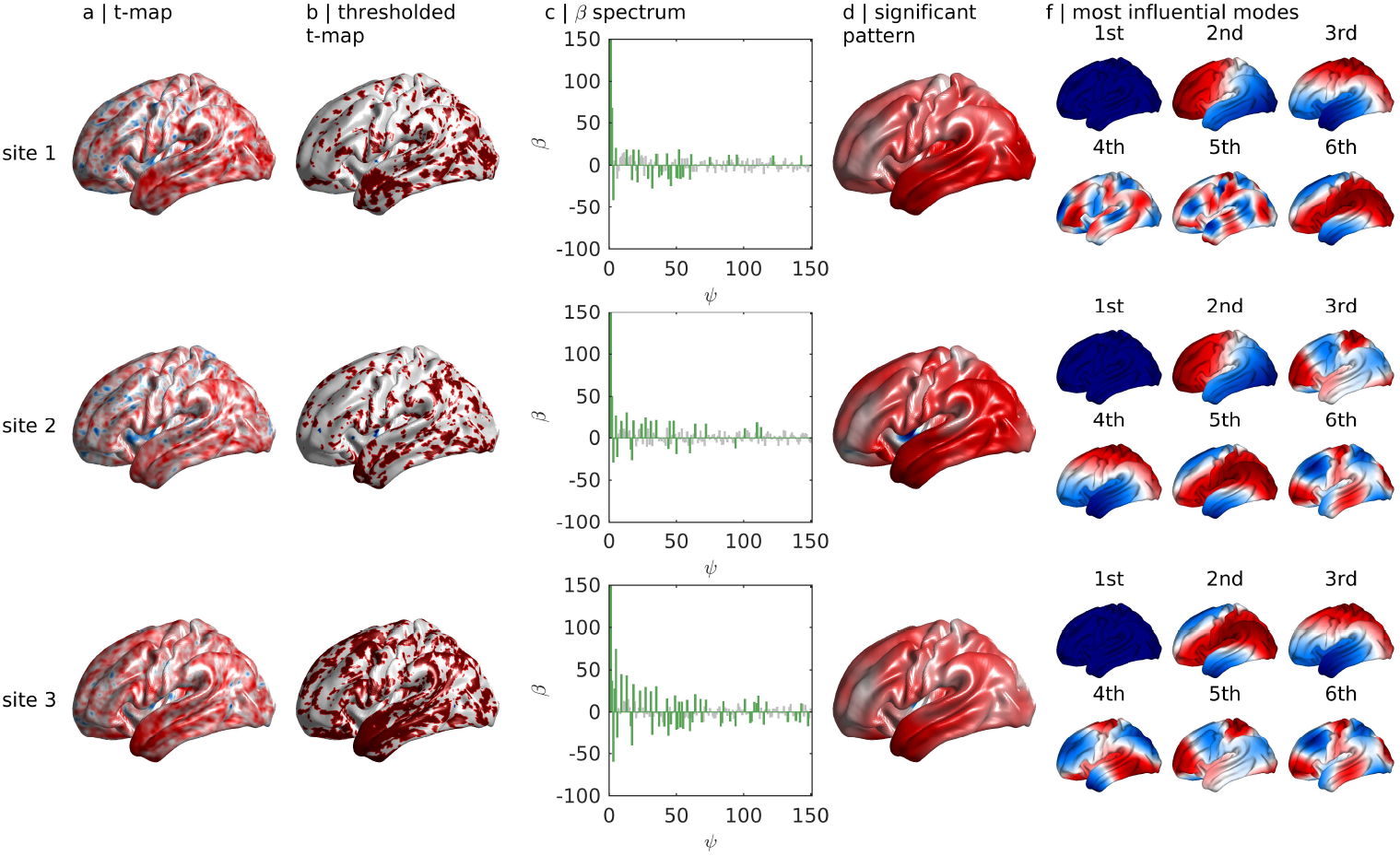
MBM analyses of CT differences between patients with Alzheimer’s disease and healthy controls across three sites. (a) Unthresholded *t*-maps of the three sites. Negative–zero–positive values are colored as blue–white–red, with positive values indicating reduced thickness in patients. (b) Thresholded *t*-map at *p*_unc_ ⩽ 0.05. Red and blue denote significantly thinner CT in patients and controls, respectively. (c) *β* spectrum of the unthresholded *t*-map. The *β*’s of the significant modes, obtained via permutation testing at *p*_unc_ ⩽ 0.05, are colored green. (d) Significant pattern obtained by combining the significant modes weighted by their *β*s. (e) Six most influential modes, i.e., significant modes with largest absolute *β* values. The signs of modes with negative *β*’s were flipped to better visualize the similarity between the modes and the significant patterns. The number denotes the order of influence, not the mode index.

Figures 10a and b show the effect of spatial smoothing on the unthresholded and thresholded *t*-maps obtained for each of the three sites with FWHM = 0, 10, 20, 30 mm, respectively (see Section 2.4.2). Figure 10c shows the absolute values of *β* spectra of the unsmoothed data for each of the three sites obtained with MBM. Here, we consider *N* = 500 modes to observe the results across a broad range of spatial scales ranging from 18 mm and larger and to more comprehensively evaluate the spatial frequency content of the data, as outlined below. The statistically significant *β*’s are shown by the green bars. For SBM, smoothing generally increases the consistency of the spatial maps observed across sites, such that nearly the entire brain shows a CT reduction in patients compared to controls at *p*_unc_ < 0.05 (Fig. 10b). The MBM *β* spectra also offer qualitative evidence for consistency across sites, with similar modes being identified as significant, particularly at lower spatial frequencies. Figure 10d shows the proportion of significant modes in each eigengroup. These proportions are high in coarse-scale, long-wavelength eigengroups, particularly groups 0 to 7, corresponding to modes 1 to 64 and wavelengths ⩾ 51 mm. Moreover, the first, global mode is also always significant, which is not observed in the analysis of sex differences (Fig. 7). Thus, while Alzheimer’s disease is associated with global reductions in CT, sex differences tend to be more focal.

**Figure 10.**
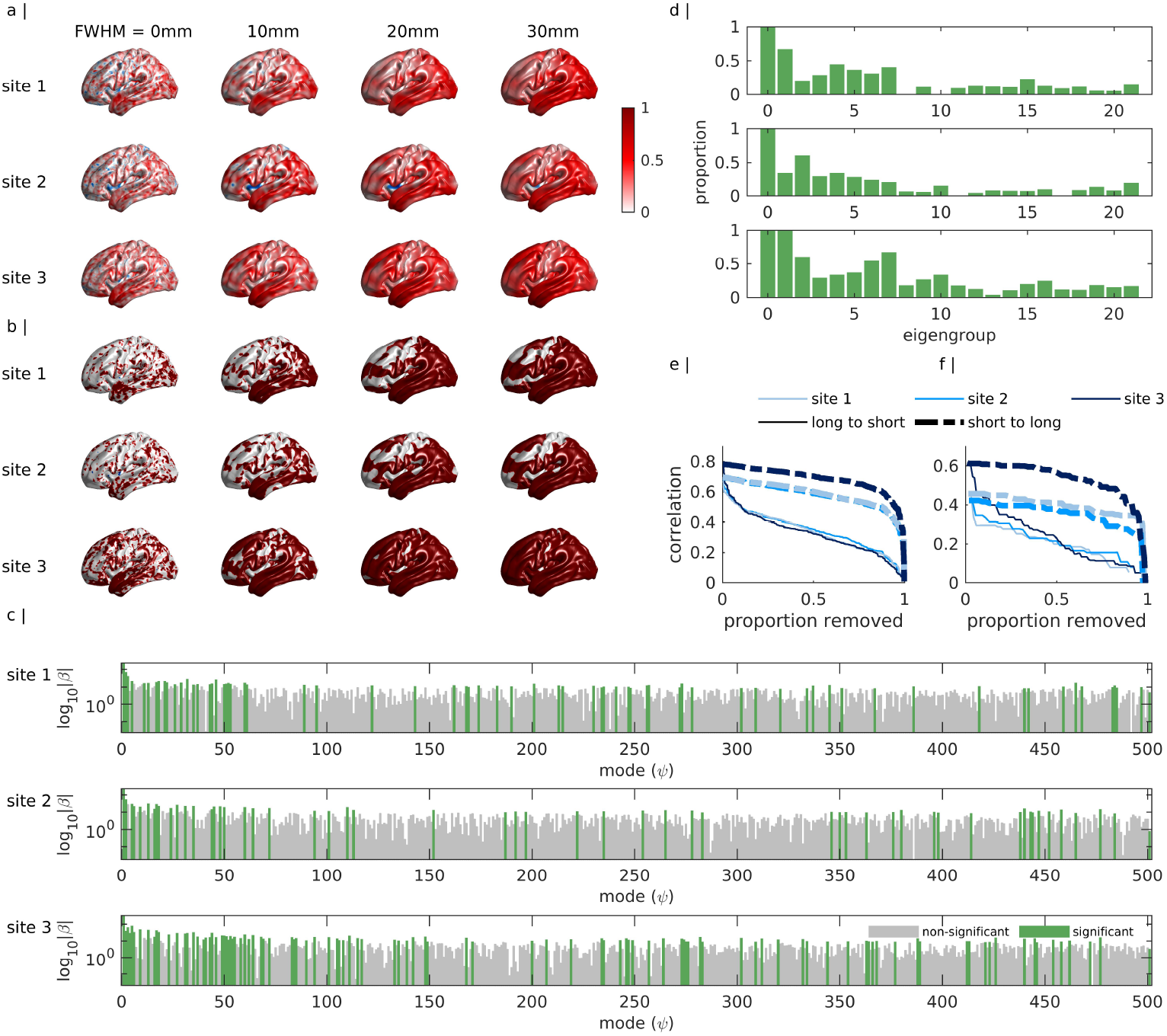
Comparing the performance of SBM and MBM analyses of CT differences between patients with Alzheimer’s disease and healthy controls across three sites. (a) Unthresholded *t*-maps of the three sites in the Alzheimer’s study with FWHM = 0, 10, 20, 30 mm. Negative–zero–positive values are colored as blue–white–red, with positive values indicating reduced thickness in patients. (b) Thresholded *t*-maps (*p*_unc_ ⩽ 0.05) of the three sites with FWHM = 0, 10, 20, 30 mm. Red and blue denote significantly thinner CT in patients and controls, respectively. (c) Absolute values of *β* spectra of the three sites without smoothing in log scale. Green and gray bars show significant and not significant *β*s, respectively. (d) The proportion of significant modes in each approximate eigengroup. (e) Correlation between the empirical *t*-map and its mode-derived reconstruction obtained using the full *β* spectrum, after removing modes in order of decreasing or increasing spatial wavelength. (f) Correlation between the empirical *t*-map and its mode-derived reconstruction obtained using only significant modes from the full *β* spectrum, after removing modes in order of decreasing or increasing spatial wavelength.

To evaluate the spatial frequency content of the difference maps in more detail, Figs. 10e and f show how correlations between the empirical and reconstructed *t*-maps are affected by the removal of a proportion of eigenmodes from the reconstruction (see Section 2.5). The reconstruction uses modes from the full *β* spectrum in Fig. 10e and from the *β* spectrum of significant modes in Fig. 10f. The correlation declines more rapidly when removing long-wavelength modes first, indicating that they make a dominant contribution to CT differences. These findings indicate that CT differences in Alzheimer’s disease are preferentially expressed over coarse scales spanning nearly the entire brain. Such broad patterns will be missed by classical analyses that focus only on point-wise inferences.

Cross-site consistency is more directly quantified in Fig. 11. Here, we considered *N* = 150 modes to be consistent with the other analyses in the paper and also because the group differences are primarily expressed over coarse scales, as observed in Fig. 10d. Figure 11a shows the pairwise site correlations for unthresholded *t*-maps and *β* spectra, while Fig. 11b shows the binary pairwise site correlations for the thresholded *t*-maps and *β* spectra. Analysis of the unthresholded maps reveals that MBM provides highly consistent results, which are unaffected by spatial smoothing. Spatial smoothing improves the performance of SBM, but it never reaches the level of MBM.

**Figure 11.**
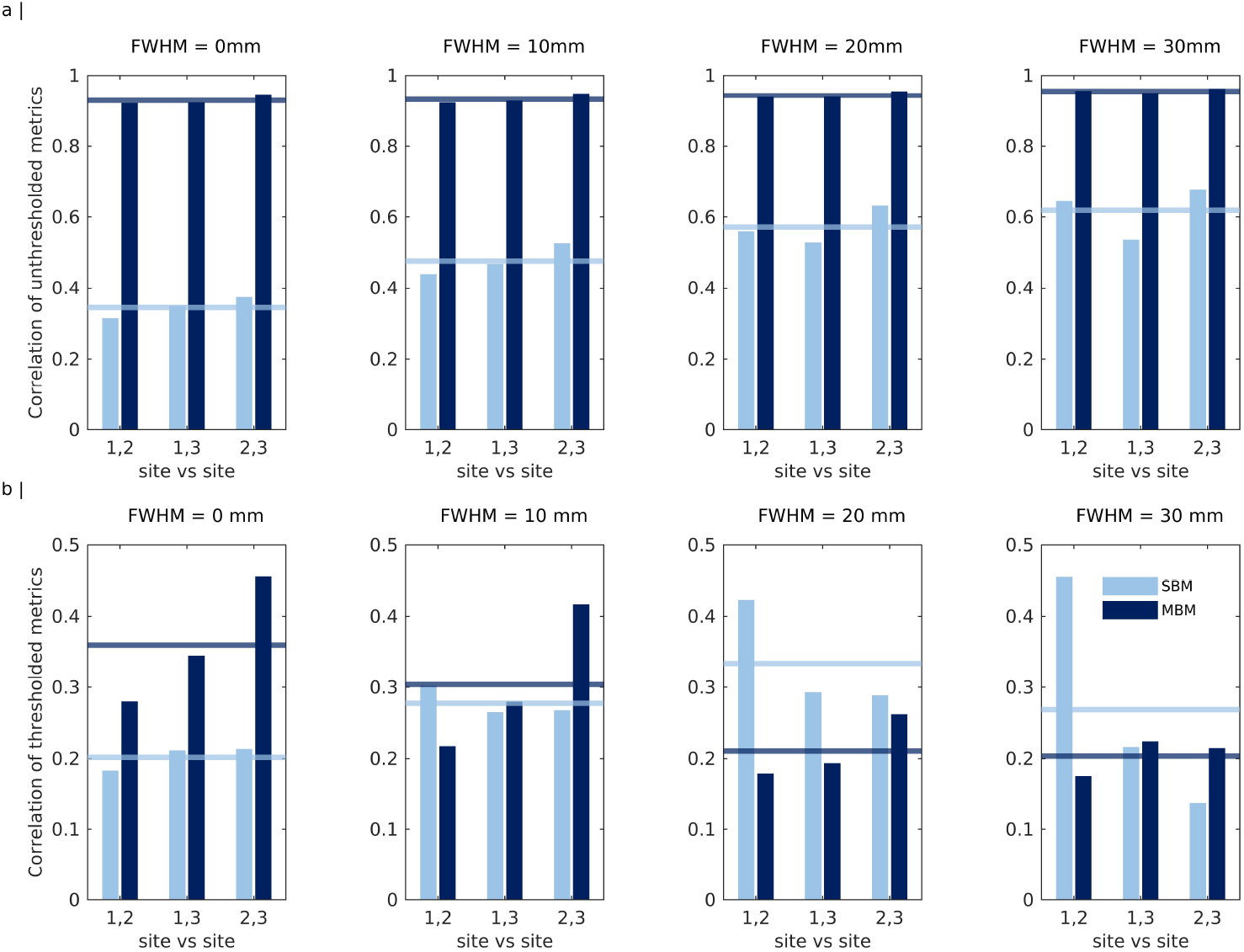
Consistency of SBM and MBM results in explaining multi-site CT differences between patients with Alzheimer’s disease and healthy controls at different smoothing kernels. (a) Pairwise correlations between sites for unthresholded results. The light and dark blue horizontal lines represent the mean correlation of the unthresholded *t*-maps (for SBM) and of *β* spectra (for MBM), respectively. (b) Pairwise binary correlations between sites for thresholded results. The light and dark blue horizontal lines represent the mean binary correlation of the thresholded *t*-maps (for SBM) and of *β* spectra (for MBM), respectively.

Considering the thresholded results, smoothing improves the performance of SBM and appears to be optimal at FWHM = 20 mm, where the average pairwise site correlations of SBM are clearly superior to MBM. However, this average SBM correlation at FWHM = 20 mm does not surpass the average MBM correlation on unsmoothed data (i.e., Fig. 11b, FWHM = 0), indicating that MBM generally yields more consistent conclusions about group CT differences in Alzheimer’s disease than classical SBM. Together, these findings support the reliability, parsimony, and simplicity of MBM given that (i) it yields more consistent inferences; (ii) it can adequately summarize the effects of interest with ∼ 150 values (i.e., the *β* spectrum) rather than the > 30, 000 vertices per hemisphere typically used in SBM; and (iii) it can be applied directly to unsmoothed data, obviating the need to select a particular smoothing kernel size and shape.

## 4. Discussion

In this paper, we developed a method for mapping neuroanatomical group differences at multiple spatial scales using a basis set of eigenmodes derived from cortical geometry, called MBM. Our analysis shows that, when compared with classical SBM, MBM shows comparable or superior performance in terms of both accuracy—i.e., capturing ground-truth differences in simulated data—and consistency—i.e., yielding consistent findings under sampling variability in both simulated and empirical scenarios. MBM also enables a spectral decomposition of the group differences, offering insights into the spatial scales at which those differences are most salient. In contrast, SBM is restricted to a single scale that is defined by the vertex mesh resolution and smoothing kernel applied. MBM does not require a choice of a specific smoothing kernel. Furthermore, MBM offers the added advantage of directly linking group differences to an underlying generative process, where the differences are modeled as resulting from the differential involvement of distinct resonant modes of brain structure.

### 4.1. MBM yields more accurate and consistent inferences about ground-truth simulations

We compared the accuracy and consistency of MBM and SBM with respect to simulated and empirical data. Our simulations offered insight into the relative performance of the two approaches under varying ratios of phenotype and (structured and unstructured) noise. This framework allowed us to evaluate the accuracy with which SBM and MBM can uncover ground-truth differences. Our analysis showed that MBM was more accurate than SBM for most simulation parameter values. The performance of MBM was particularly strong in parameter regimes associated with high levels of unstructured (i.e., Gaussian) noise, indicating that it is likely to perform better in noisy data due to the low-pass spatial filtering effect of the eigenmode reconstructions.

When structured noise dominated the ground-truth difference, MBM and SBM showed comparable mean accuracy, but MBM correlations were more variable across experiments, indicating less consistency. To the extent that structured noise in our model emulates subject-specific neuroanatomical features, this finding indicates that MBM will perform inconsistently in cases where the contribution of such features (i.e., inter-individual variability) match or swamp the ground-truth phenotype that is common to all subjects in the same group. This is because MBM is particularly sensitive to detecting structured (i.e., autocorrelated) patterns in the spatial maps. However, group mean comparisons of CT (or other neuroanatomical properties) are not likely to be very meaningful in such scenarios, as the small contribution of the ground-truth difference indicates that there is little in common between different people assigned to the same group.

Spatially smoothing the data improved the performance of SBM to the point where it matched MBM for smoothing kernels with FWHM ⩾20 mm. However, there are no consistent heuristics for choosing a proper smoothing kernel. In addition, smoothing kernels should be chosen carefully since they can impose geometric effects on the data as the kernel replaces the value at each point with a weighted average of its spatial neighbors [32]. A particular advantage of MBM is that it does not require the selection of a specific smoothing kernel size.

### 4.2. MBM yields more consistent results in empirical data

It was not possible to evaluate the accuracy of MBM and SBM in empirical data due to the lack of ground truth, but we could evaluate the consistency of the findings across multiple repetitions of the same experiment. For both the analyses of differences between sexes and between healthy controls and patients with Alzheimer’s disease, MBM was more consistent than SBM at low levels of smoothing, with the two converging at higher levels of smoothing.

In the case of sex differences, the distributions of MBM correlations were generally wider than those of SBM correlations. When interpreted with respect to our simulations, this result suggests that while there may be some consistent sex differences in CT that can be observed under resampling, they are likely subtle relative to the effects of individual variability. This interpretation aligns with ongoing debates over the consistency of sex differences in neuroanatomy [3, 18, 19].

Alzheimer’s disease is likely to be associated with a more salient and robust CT phenotype, given the well-described stages of atrophy that are known to occur in the condition [59, 60, 61]. Accordingly, both unthresholded and thresholded analyses were more consistent for MBM compared to SBM. Increasing smoothing kernel size improved the performance of SBM, but the average consistency never surpassed that observed for MBM in the unsmoothed data. These findings support the utility of applying MBM to minimally smoothed data.

### 4.3. MBM offers insights into scale-dependent group differences

Structural eigenmodes are ordered by spatial wavelength, opening the opportunity to analyze the spatial frequency content of CT differences, much like a Fourier decomposition is routinely used to examine spectral properties of EEG signals. We presented an analysis of the frequency content of CT differences in Alzheimer’s disease, showing that the proportion of significant modes was higher in eigengroups with long wavelengths, with most of the differences found for modes with wavelengths ⩾ 51 mm. The first, global mode was consistently significant across the three sites, indicating a robust global difference in CT between cases and controls. Reconstruction accuracy also declined more rapidly when removing long-wavelength modes, indicating that they make a dominant contribution to CT differences between groups. Together, these findings suggest that CT differences in Alzheimer’s disease are most salient at coarse spatial scales that are not adequately captured by classical point-wise analysis, such as SBM, where a specific resolution scale is imposed by the mesh resolution and smoothing kernel size. In contrast, MBM provides a natural way of characterizing group differences across a wide range of spatial scales.

A further consideration is that statistical inference in SBM often assumes that each point-wise location (e.g., surface vertex) is independent. This assumption is incorrect, since CT and many other neuroanatomical properties are spatially autocorrelated; i.e., the value at one point depends on others. The violation of this assumption is worsened when spatial smoothing is applied to the data. Dependencies are sometimes later considered if some form of cluster-based thresholding is applied [62], but this is an ad hoc characterization. In MBM, distinct spatial locations are not considered to be independent but instead form part of brain-wide modes with varying spatial wavelengths. Critically, since the eigenmodes themselves are orthogonal, mode-specific inference is entirely justified.

### 4.4. MBM ties neuroanatomical differences to a generative process

Classical approaches to mapping neuroanatomical differences are purely phenomenological, relying on statistical analyses to identify differences between groups without offering a direct explanation for the mechanisms through which those differences have emerged. An important advantage of MBM is that the results can be linked to a direct physical interpretation, in which group differences in neuroanatomy are explained as the involvement of different resonant modes of brain anatomy. An intuitive analogy can be drawn from plucking a violin string [31]. The eigenmodes correspond to the string’s harmonics, each of which is associated with its own preferred vibration frequency. The musical note generated by plucking the string results from a superposition of these modes. This basic idea has been used to understand how the structure of a system constrains its dynamics in diverse areas of physics and engineering, including the electromagnetic response of different media [20], the vibrational patterns of different structures [21], and aeroelasticity [22].

In the cortex, these resonant modes define the principal axes of structural variation and thus represent a fundamental basis set for understanding anatomical constraints on any spatially patterned process. For instance, the second, third, and fourth eigenmodes considered here correspond to spatial variations along the anterior-posterior, dorso-ventral, and medial-lateral axes, which are known to define many fundamental properties of cortical organization, such as regional variations in cell density [63], CT [64], genetic influences on neuroanatomy [65], and gene expression gradients that shape brain development [66, 67].

An important choice in such analyses concerns the neuroanatomical properties that should be used to define the eigenmodes. Two distinct approaches have emerged in the literature. One approach involves deriving eigenmodes from a discrete, graph-based model of the connectome, under the assumption that inter-regional connectivity represents the primary anatomical constraint on dynamical processes in the brain [23]. The other approach involves deriving eigenmodes from a model of the geometry of the cortex, as used here. This approach follows from a specific form of neural field theory, a well-validated biophysical model of brain dynamics that characterizes how dynamical processes spatially propagate as waves through a continuous cortical medium [46]. Recent work indicates that geometric modes offer a more parsimonious account of diverse aspects of brain function than connectome eigenmodes [15]. Geometric eigenmodes also offer a practical advantage, as they can be extracted from T1-weighted MRI data alone using standard procedures, whereas connectome eigenmodes rely on the application of complex preprocessing pipelines to diffusion MRI data, which require many choices that can affect the final results [68, 69, 70]. For these reasons, we have focused on geometric modes in our analysis, but the approach developed here is sufficiently general that it can be used with any anatomical basis set.

## 5. Limitations and Conclusions

We derived the geometric eigenmodes using a population-average template surface, which does not completely account for individual differences in brain shape. While low-frequency modes tend to be consistent between people [41], the spatial patterns of high-frequency modes tend to diverge due to individual differences in neuroanatomy. This divergence makes it difficult to compare results across individuals. Past work suggests that modes derived from a population-average template can reconstruct brain function to a degree that is comparable to individual-specific modes [15], but further work is required to develop techniques that can better capture individual differences in eigenmode architecture. Future work could also incorporate volumetric analyses in subcortical regions to enable whole-brain inferences, given the strong coupling between geometry and function found in these areas [15].

In summary, we have introduced here a new multiscale approach, which we call mode-based morphometry (MBM), for mapping neuroanatomical differences between groups where the differences are modeled as arising from the involvement of distinct, resonant modes of brain anatomy. Using both simulated and empirical data, we show that MBM offers more accurate and consistent inferences than classical approaches (i.e., surface-based morphometry), while also providing insights into the spatial frequency content of the differences and allowing a direct link to putative generative physical processes.

## Data and code availability

Raw and preprocessed HCP data can be accessed at https://db.humanconnectome.org/. Raw and preprocessed OASIS-3 data can be accessed at https://www.oasis-brains.org/. An open-source toolbox implementing MBM will be available at https://github.com/BMHLab/MBM upon publication of the article. Code and sample data to reproduce the analysis results and figures of this study will be openly available at https://github.com/BMHLab/MBM paper upon publication of the article.

## Acknowledgments

Data were provided in part by the Human Connectome Project, WU-Minn Consortium (Principal Investigators: David Van Essen and Kamil Ugurbil; 1U54MH091657) funded by the 16 NIH Institutes and Centers that support the NIH Blueprint for Neuroscience Research; and by the McDonnell Center for Systems Neuroscience at Washington University. Data were provided in part by OASIS: Longitudinal Multimodal Neuroimaging: Principal Investigators: T. Benzinger, D. Marcus, J. Morris; NIH P30 AG066444, P50 AG00561, P30 NS09857781, P01 AG026276, P01 AG003991, R01 AG043434, UL1 TR000448, R01 EB009352. AV-45 doses were provided by Avid Radiopharmaceuticals, a wholly owned subsidiary of Eli Lilly. This work was supported by the MASSIVE HPC facility (www.massive.org.au) [71]. AF was supported by the Sylvia and Charles Viertel Foundation, National Health and Medical Research Council (IDs: 1146292 and 1197431), and Australian Research Council (IDs: DP200103509 and FL220100184)

