## Supplementary for "Mode-based morphometry: A multiscale approach to mapping human neuroanatomy"

Table S1: Wavelengths and eigenmode membership for each eigengroup on a sphere of $R_s$ = 67 mm.

| Eigengroup | Wavelength (mm) | Eigenmodes included in the eigengroup |
|---|---|---|
| 0 | - | 1 |
| 1 | 297.7 | 2–4 |
| 2 | 171.9 | 5–9 |
| 3 | 121.5 | 10–16 |
| 4 | 94.1 | 17–25 |
| 5 | 76.9 | 26–36 |
| 6 | 65.0 | 37–49 |
| 7 | 56.3 | 50–64 |
| 8 | 49.6 | 65–81 |
| 9 | 44.4 | 82–100 |
| 10 | 40.1 | 101–121 |
| 11 | 36.6 | 122–144 |
| 12 | 33.7 | 145–169 |
| 13 | 31.2 | 170–196 |
| 14 | 29.1 | 197–225 |

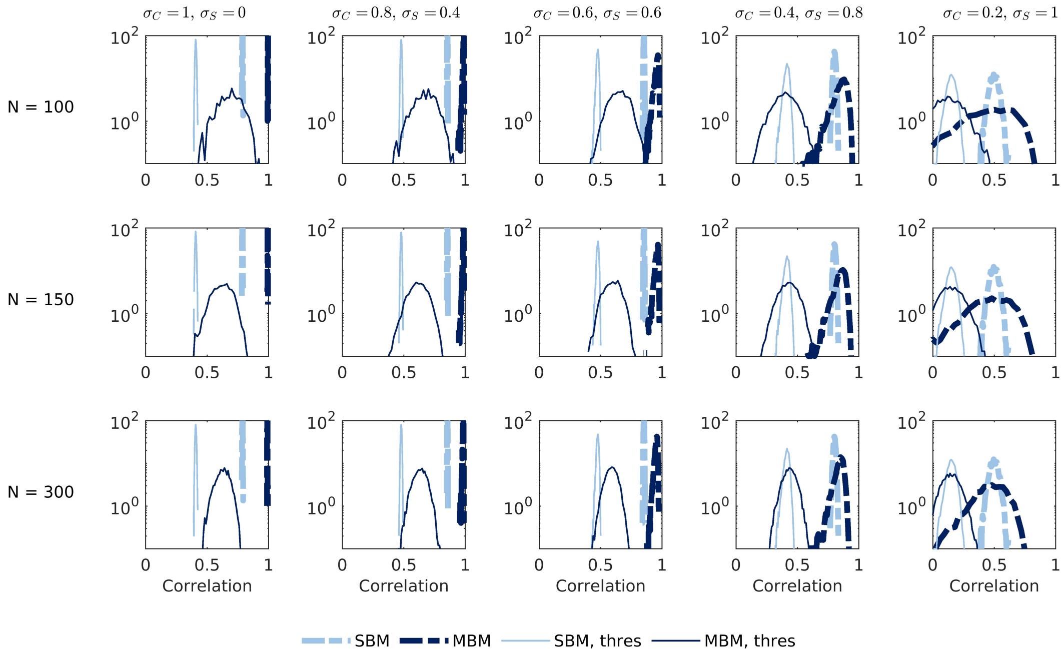


Figure S1: Distribution (in log scale) of pairwise correlations between experiments in the realistic parameter regimes and for different numbers of eigenmodes, $N$. The panels show correlations for experimental $t$-maps (SBM), correlations for experimental $\beta$ spectra (MBM), binary correlations for thresholded $t$-maps (SBM, thres), and binary correlations for statistically significant $\beta$ spectra (MBM, thres).

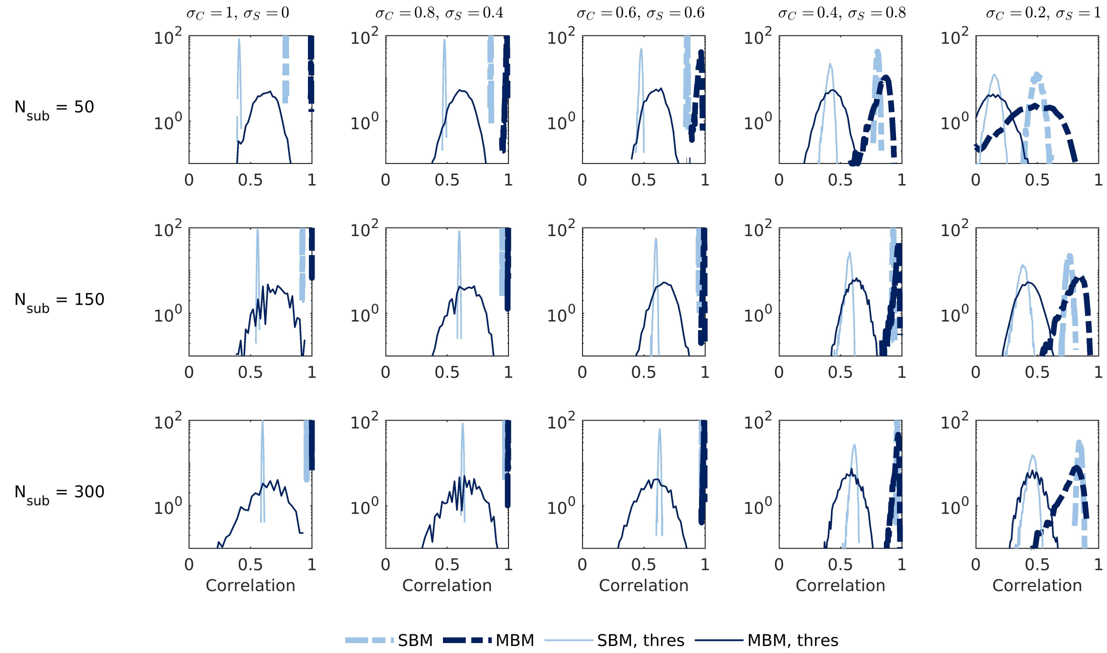


Figure S2: Distribution (in log scale) of pairwise correlations between experiments in the realistic parameter regimes and for different numbers of subjects, $N_{sub}$. The panels show correlations for experimental $t$-maps (SBM), correlations for experimental $\beta$ spectra (MBM), binary correlations for thresholded $t$-maps (SBM, thres), and binary correlations for statistically significant $\beta$ spectra (MBM, thres).

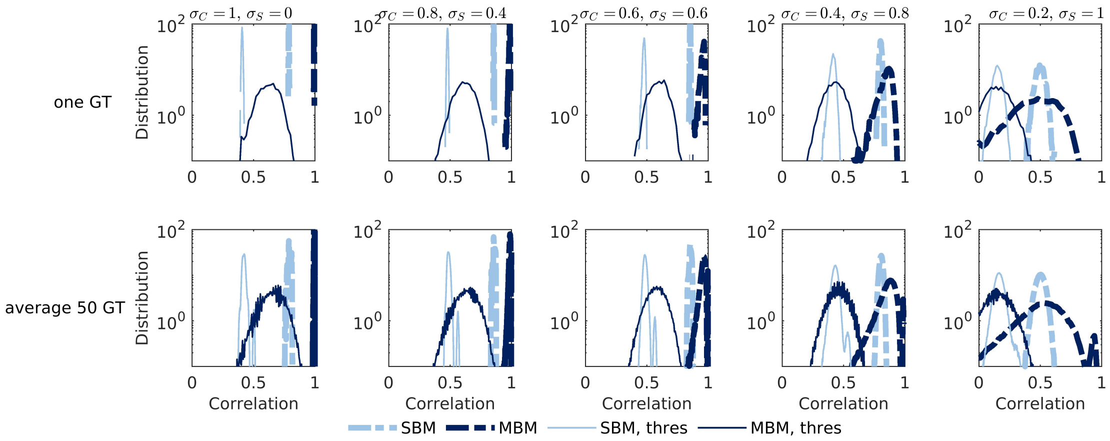


Figure S3: Distribution (in log scale) of pairwise correlations between experiments in the realistic parameter regimes and for different ground truths (GT). The panels show correlations for experimental $t$-maps (SBM), correlations for experimental $\beta$ spectra (MBM), binary correlations for thresholded $t$-maps (SBM, thres), and binary correlations for statistically significant $\beta$ spectra (MBM, thres).

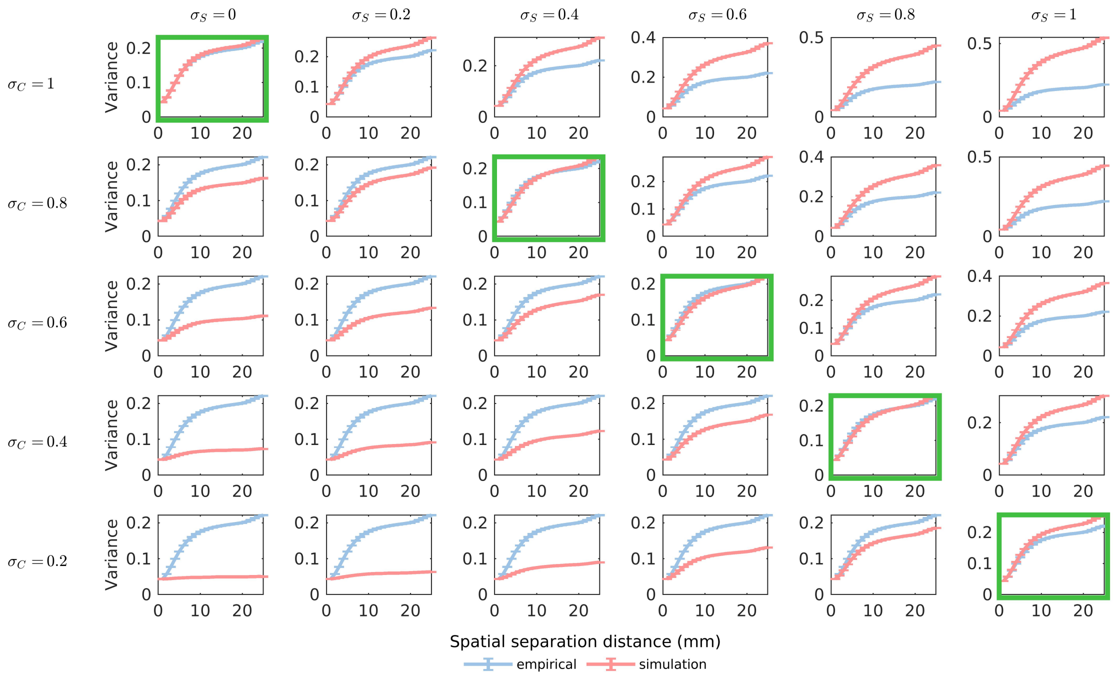


Figure S4: Mean variograms (after subtracting the minimum offset) of empirical and simulated maps for combinations of $\sigma_C$ and $\sigma_S$. The green boxes highlight the realistic regimes where the generated maps have a similar spatial structure as the empirical data (distance $\approx 0$).

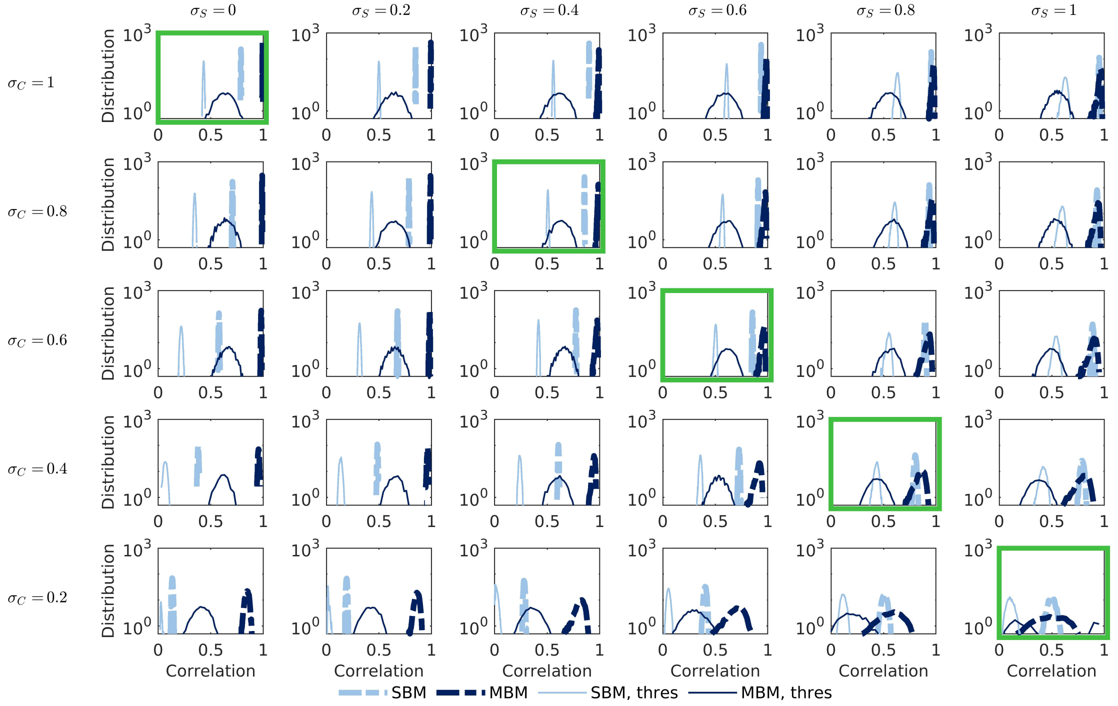


Figure S5: Distribution (in log scale) of pairwise correlations between experiments for different combinations of $\sigma_C$ and $\sigma_S$ with FDR correction. The panels show correlations for experimental $t$-maps (SBM), correlations for experimental $\beta$ spectra (MBM), binary correlations for thresholded $t$-maps (SBM, thres), and binary correlations for statistically significant $\beta$ spectra (MBM, thres).
